# Perceived Stress in First-Year Medical Students and its Effect on Gut Microbiota

**DOI:** 10.1101/854174

**Authors:** Matthew Revere Rusling, Joseph Johnson, Aaron Shoskes, Chunfa Jie, Li-Lian Yuan

## Abstract

Medical students are constantly under stress caused by strenuous medical programs, which may exert persistent physical and psychological effects on their well-being. Using medical students as a model population, this work explores the gut microbiome as a potential contributing mechanism for why individuals exposed to similar stimuli react variably. We evaluated the relationship of gut microbiome composition of first year medical students and stress resilience over a period of 4 months. Our objective was to identify gut microbiome characteristics of individuals that showed long-term stress resilience. Students were voluntarily recruited and screened for lifestyle and environmental factors at 3 timepoints during the first semester. Fecal samples were also collected at each timepoint. In order to identify candidates with stress resilience, their perceived stress and depression levels were normalized and summed to produce a psychologic index score. The most notable finding is a correlation between psychologic resiliency of Bacteriodete:Firmicute abundance as well as a relationship between durable resiliency and microbiome stability. Phylogenetic assembly of participants by microbiome relatedness found that 100% of subjects who were resilient to stress across *all* timepoints (n=8) were phylogenetically clustered in adjacent positions, showing a high degree of temporal stability. Of participants who were not durably resilient to stress, only 62% of participants (n=8) showed microbiomes that were phylogenetically related across the same 4 month period. We identified 2,102 Operational Taxonomic Units (OTUs) which were unique to the durable resilience group and 94 OTUs which were unique to the susceptible group. Of the 4,794 observed OTUs, 6.1% (n=294) were significantly different between groups. These findings support that the gut microbiome may play an important role in stress resilience at a time scale of 4 months. A better understanding of the role of the gut microbiome in stress resilience may shed light on potential treatment to reduce stress/anxiety in general, as well as to promote wellbeing of our future health care providers and physicians.

## Introduction

The scientific community has increasing interest in the gut microbiome as the scope of the host functions that it serves have become better understood. Mechanistic exploration of the role of the gut microbiome on host function has revealed that the microbiome exerts potent effects on its host ranging from modulating brain barrier permeability (Braniste et al., 2014) to contributing to metabolic syndrome and obesity (Tilg and Kaser, 2011). Some investigators have calculated that the gut microbiome contains “100 trillion microorganisms from 40,000 species with 100 times the number of genes [as found] in the human genome.”(Mayer, 2011). As a result of evolution from a common ancestor and synergistic co-evolution between multicellular and unicellular organisms throughout biologic history, our neuroactive substrates are also utilized by bacteria for cell-cell signaling (Mayer, 2011). The profile of neuroactive substrates produced by the gut microbiome is becoming well characterized and includes serotonin, short chain fatty acids, GABA, dopamine and norepinephrine (Hyland and Cryan, 2010; Mayer, 2011). Within the human host, through interactions with the myenteric nervous plexus, microbiome neurotransmitter production by the proximal gut essentially provides bottom-up visceral sensory feedback from the microbiome to the brain via Cranial Nerve X and other projections (Hyland and Cryan, 2010; Mayer, 2011). As a known example of bottom-up feedback at acute timescales, food consumption has been found to downregulate pain perception. One mechanism that has been proposed for this phenomenon is that consumption of food causes increased microbiome neurotransmitter production which increases feedback onto neuroendocrine cells. Increased neuroendocrine cell stimulation outflows to lamina I of the spinal cord, ultimately synapsing to the central monoaminergic nuclei of our neural centers (Mayer, 2011). It is through modulation of these ascending pathways which are closely associated with pain pathways that increased luminal microbiome activity is argued to downregulate pain perception in mammals (Mayer, 2011). Changes to microbiome taxonomic composition have the downstream effect of altering the translational profile of microbiome products (Karl et al., 2018). Thus, when taxonomic changes occur, regulatory feedback from the gut lumen to the myenteric plexus is altered and ultimately projections to the central nervous system are up or downregulated (Hyland and Cryan, 2010).

Additionally, prior research has identified that the gut microbiome is involved in the development of at least some psychiatric conditions such as anxiety, depression, schizophrenia and bipolar disorder (Kelly et al., 2015; Bastiaanssen et al., 2019). Recognizing the multi-faceted role that the gut microbiome plays in host physiology and mental illness, we sought to explore the role of the gut microbiome in human stress resiliency. Resiliency can be considered as an individual’s total capacity to minimize their maladaptive cognitive response (such as depression, or anxiety) to a novel stressor. While much attention has been paid to the psychology of stress resilience and successful thought patterns which contribute to resilience, we are only scratching the surface of understanding the mechanistic biology underlying resilience and why some individuals are more resilient than others.

This work is a longitudinal study which follows 40 medical students through the first 8 months of their curriculum to retrospectively identify microbiome features of stress resilient and stress susceptible students as measured by serum cortisol and validated self-reported stress and depression scores. Our goal was to characterize conserved features of the gut microbiome of stress resilient individuals. Recognizing that the perception of stress involves psychologic and physiologic components, we hypothesize that changes to the composition of the gut microbiome may be related to stress intolerance.

Evaluation of medical student microbiomes and stress responses present a particularly well-suited opportunity to explore this question (Dyrbye et al., 2008) (Tateno et al., 2018). Medical school is a notoriously stressful academic setting in which major stressors such as written and practical exams are applied at the same timepoints across a sizeable cohort (ranging from 60 to 250 students). These students have also moved from different parts of the country to the same location at roughly the same time, drink from the same water sources and consume foods from more or less the same markets; making them an ideal population for the study of the gut microbiome and stress resilience. Finally, among these medical students why is it that some are resilient to comparable stressors while others experience marked involuntary increases in cortisol, stress and depression as a maladaptive response to the novel rigorous eustress of their medical education?

## Methods

### Recruitment and selection

First and second year medical students at Des Moines University (DMU) were voluntarily recruited to this study through their class social media pages. Project protocols and procedures were approved by DMU Institutional Research Board (IRB). When considering applicants, our inclusion criteria were: enrollment at DMU’s medical school, age 21-41. Our exclusion criteria were: extreme diet (such as paleo, vegan, keto), antibiotic usage in past 3 months, probiotic use, history of anxiety, current medication use for anxiety or depression, pregnancy, obesity, competitive athlete, current tobacco use, history of chronic allergy, immunosuppressed illness, Irritable bowel disease or syndrome, GERD with PPI use, or weight under 50 kg. We recruited 31 students who met our criteria and were willing to provide psychologic and microbiome data at 3 timepoints, each in August, October and December.

### Data collection

At each timepoint we collected Cohen’s Perceived Stress, PHQ-9 Depression scores, and measured serum cortisol levels in addition to administering uBiome kits for metagenome fecal sample collection. Human tissue samples were collected following a protocol approved by DMU Institutional Biosafety Committee (IBC). Serum cortisol levels were obtained by venipuncture. Results were anonymized protect participants’ identify and unique ID’s were assigned to each participant to allow tracking of participants for analysis. An approved protocol was followed for human data protection and management.

### Psychologic Index Score (PIS)

Results from Cohen’s Perceived Stress and PHQ-9 surveys were normalized against the total point value of the surveys. These were summed with a cortisol value which was normalized against the highest cortisol value observed in the dataset. Normalized values were used to produce a Resiliency Index Score (RIS). The RIS was used to categorize individuals as stress susceptible or resilient across timepoints, and also characterize samples as originating from a high or low stress participant at a discrete timepoint.

### Workup and filtering of metagenome

DNA extraction, amplification and sequencing performed by uBiome using their SmartGut® Kit. 16S V4 rRNA gene was amplified using 515F/806R primers and sequenced on NextSeq 500 platform producing 150bp paired-end reads. Reads were filtered below an average Q-score >30 and truncated at 125 bp and 124 bp for forward and reverse reads, respectively. Reads containing >8 consecutive nucleotides were discarded. Chimera removal was performed before reads were compared against version 123 of the SILVIA database, and those <77% similar to known 16s sequences were discarded. This filtered data was then imported into QIIME 1.9.0 and initially processed on that platform.

## Results

### Psychological and physiological stress response to medical school

In order to assess the level of perceived stress in first year medical students, PHQ-9 and Simpson Stress survey responses were collected from all 30 participants at 3 timepoints of the study. Among the three timepoints, August represented a time before the school started, October and December coincided with mid-term and final exams, respectively (**Figure 1A**). Owing to participant withdrawal or failed microbiome sample workup and/or sequencing, we successfully recorded 69 discrete collections including PHQ-9, serum cortisol, Simpson stress, and sequenced fecal sample. Complete data set across all three timepoints was collected from 16 out of 30 participants. Of these discrete data points, we recorded 18 and 61 instances of low and moderate perceived stress respectively (Scores: low<14, moderate=14-27), and 59, 7, and 3 instances of minimal, mild and moderate depression respectively. Due to the small sample size of participants and the moderate range of reported stress levels between high stress and low stress individuals this was not an ideal opportunity to identify specific phyla changes and broader microbiome features were instead explored.

**Figure 1.**
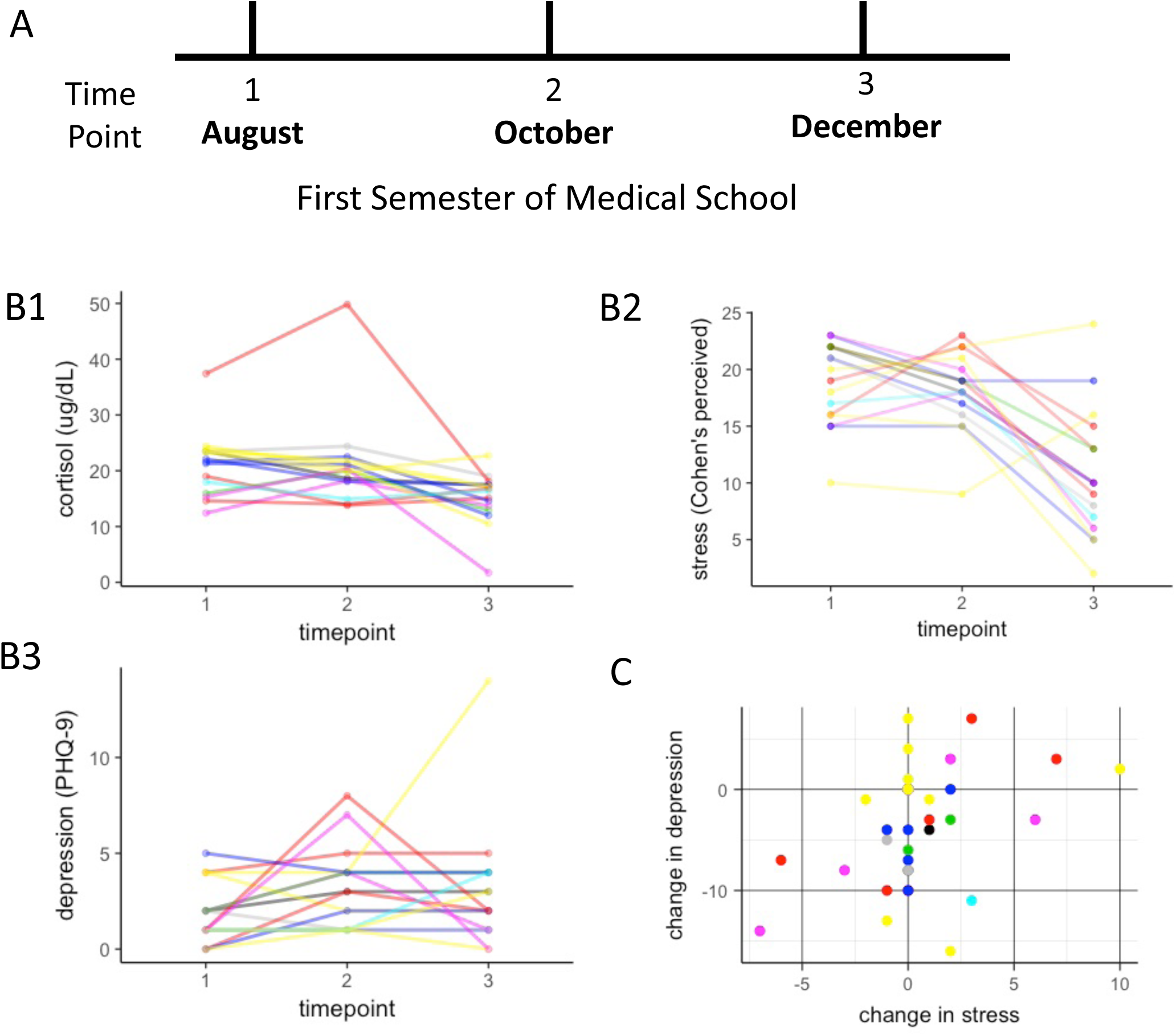
Physiological measurement and psychological survey responses indicating the stress level of first year medical students. **A.** Experimental timeline of survey responses, blood cortisol measurement, and fecal sample collection from medical students (n=30) at three time points of the first semester**. B.** Cortisol (B1) and stress (B2) levels decreased significantly (Pearson’s Correlation, p<0.01, R=-0.35 and p<0.01, R=-0.55, Respectively) while depression did not significantly change across timepoints (B3). Each color represents each individual. **C.** Scatter plot captured 51 individuals expressing moderate stress levels, 4 instances of moderately depression and 2 instances of moderately severe depression (see text for classification criteria).

Across all participants and timepoints (n=69), cortisol levels changed significantly between timepoints (**Figure 1 B1**, GLM, Wilk’s Lambda=0.42, p=0.001). Stress scores changed significantly during the experiment between timepoints (**Figure 1 B2**, GLM, Wilk’s Lambda=0.26, p<0.001). At an individual level, an increase in stress between timepoints 1 and 2 did not predict the direction of change between timepoints 2 and 3 (Pearson’s correlation, p=0.3), while a change between timepoints 1 and 2 was significantly predictive of a change in depression in the *opposite* direction at subsequent timepoints (Pearson’s correlation, p=0.04, r=-0.5). Depression scores did not significantly change during the study (**Figure 1 B3**, GLM, Wilk’s Lambda=0.845, p=0.24). Overall, physiological measurement and self-reported stress and depression perception exhibited high levels of variance and heterogeneity.

### Resilience Stress Index (RIS)

The high degree of directional inconsistency between measures across timepoints observed in Figure 1 lead us to select for an extreme phenotype of stress resiliency by creating a Resiliency Index Score (RIS). Previous studies of human resilience have used either depression or perceived stress to quantify the resilience response (Oken et al., 2015; Winslow et al., 2015). After observing non-correlated changes between perceived stress and depression we chose to combine these normalized scores to increase our specificity in identifying truly resilient individuals. The RIS used here was calculated by normalizing the PHQ-9 and Simpson’s Index scores against their total possible scores (27, 40 respectively) and summing the normalized value into a 2 point-maximum score. Cortisol was excluded from the RIS to remove for confounding effects of cortisol’s time-dependent variability and to emphasize how the gut microbiome may effect stress perception.

Across all participants regardless of their gender, RIS scores were found to decrease significantly throughout the experiment (**Figure 2**, Pearson’s correlation, r=-0.41, p<0.001). For participants with 3 timepoints of data available, and to support the use of an RIS as a method to identify an extreme stress resilience phenotype, an individual’s change in RIS score between timepoints 1 and 2 was positively correlated with a change in RIS score at the subsequent timepoint for that individual (Pearson’s correlation, p=0.003, r=0.69). Female gender was associated with a significantly higher RIS score (ANOVA, F=5.89, p=0.018).

**Figure 2.**
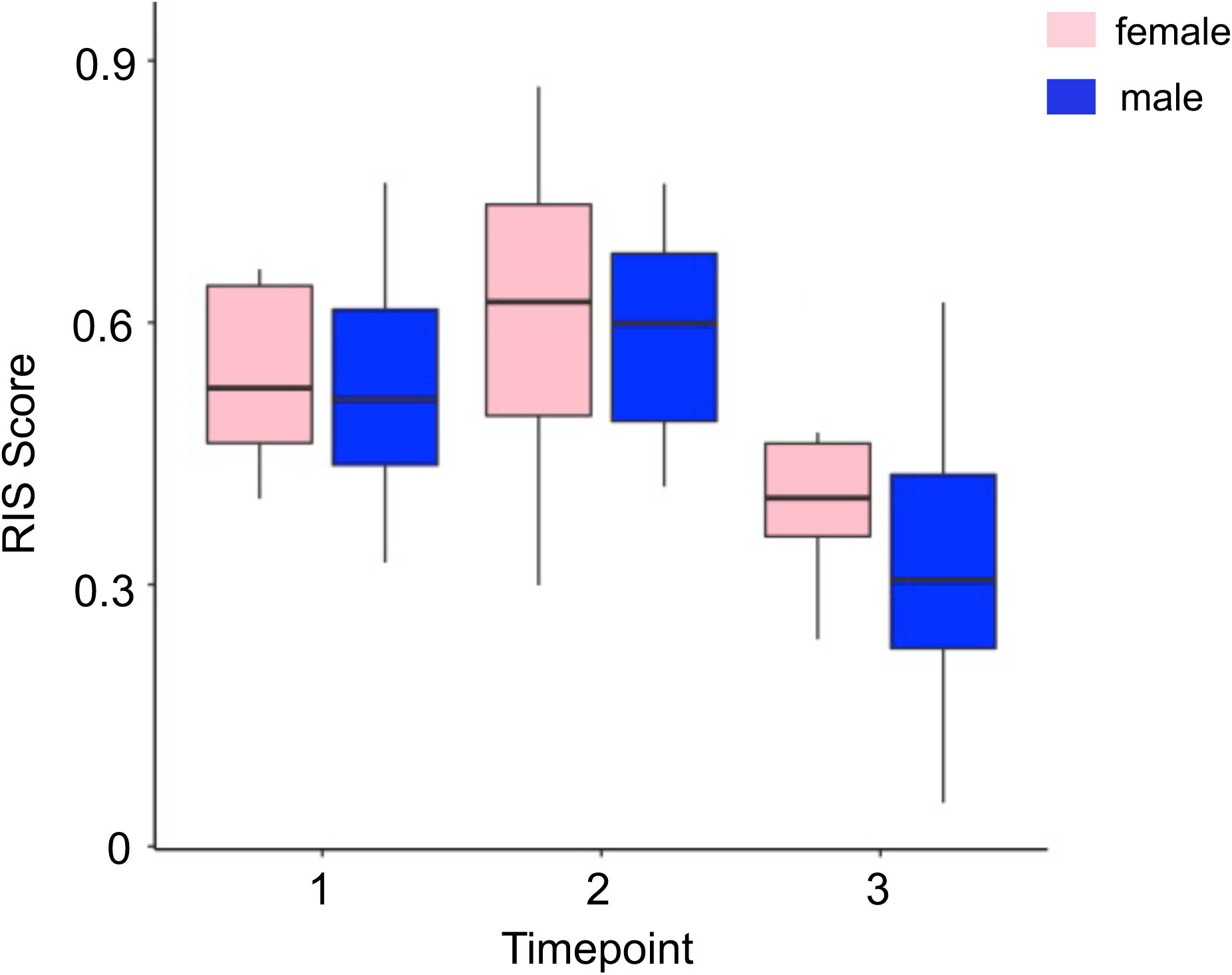
Gender and temporal profile of RSI scores. The RSI score, as a combined index of perceived stress and depression levels, decreased across time (Pearson’s correlation, r=-0.41, p<0.001). Comparison between male and female participants revealed s gender difference in RSI scores with females having significantly higher RSI scores on average (ANOVA, F=5.89, p=0.018).

Of the 16 participants who provided 3 timepoints of complete data, 8/16 of these participants showed RIS scores which either remained stable or decreased across the experiment, showing an extreme resilience phenotype (**Figure 3**, blue). The other 8 participants showed a stress susceptible phenotype, in which the RIS score increased in at least one timepoint (**Figure 3**, red). By focusing on individuals with index scores that either remained constant or decreased during the stress of medical school we were able to identify 8 participants with durable psychologic resilience. We compared them against participants whose scores increased at any period during the experiment. In addition, we sought to identify taxonomic changes which were associated with either an increase or decrease in psychologic index score between timepoints *(Pink=stress susceptible, blue=durable resistance)*.

**Figure 3:**
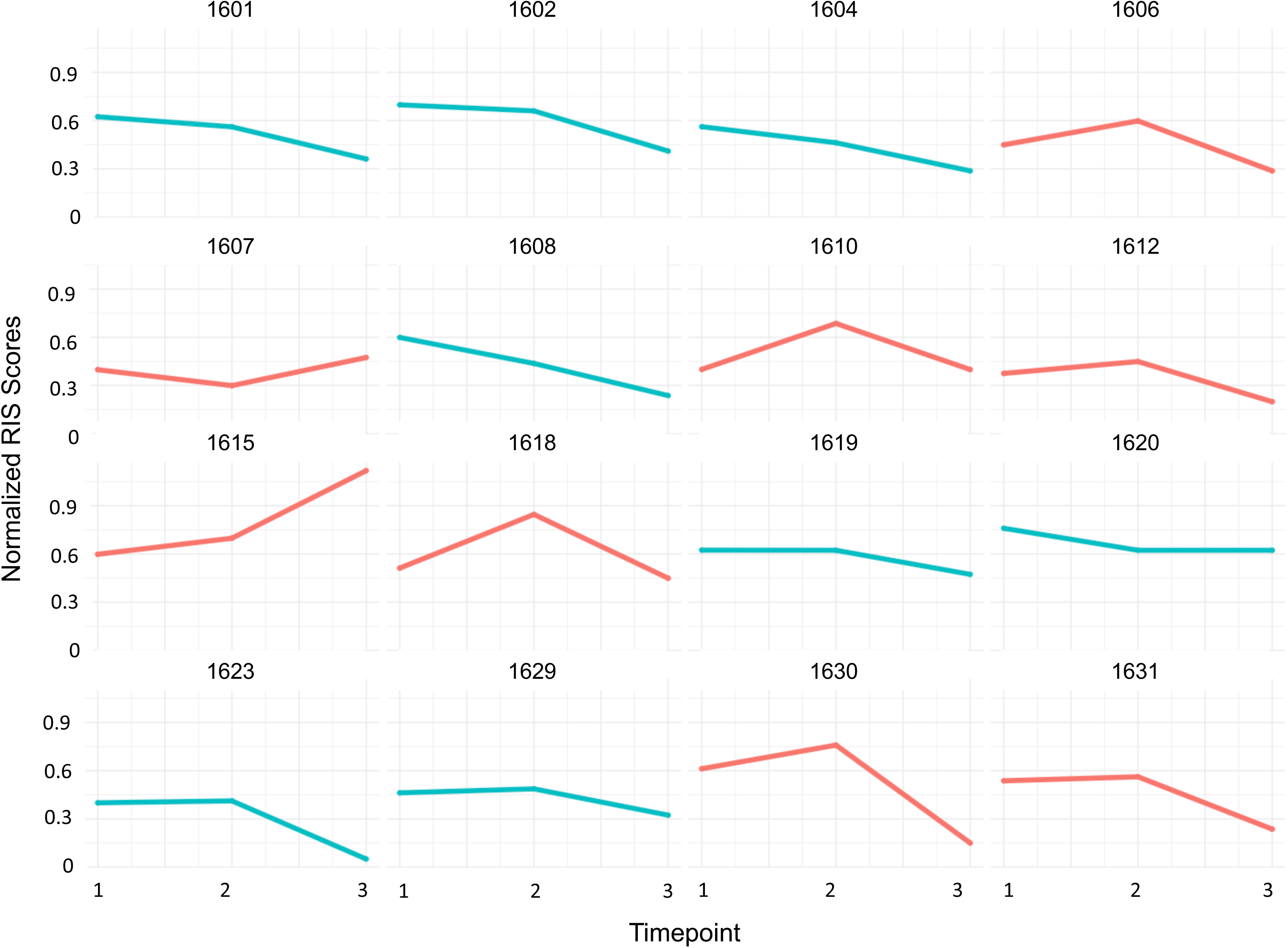
Temporal RSI profile of individual participants. A complete date set from all three time points were obtained from 16 participants (1601, 1602, 1604 … etc.). Based on their temporal profiles of RSI that showed a decrease during the first semester of medical school, a group of 8 participants was identified as stress resilient (Blue)..

### Microbiome overview

Microbiome sequencing was successful for 69 samples after filtering. 5,653 unique 97% similar Operating Taxonomic Units (OTUs) were observed, with a total of 9,754,866 sequences. The mean number of samples was 14,1394±9,7067 stdev. Alpha diversity calculations were made using a not otherwise filtered OTU table. Beta diversity comparisons were performed on OTU tables which were filtered for observations that only occurred in one sample, and for features which composed less than 0.0005% of the microbiome. To control for differences in total sequencing numbers, OTU tables were normalized. After these filtering step prior to beta diversity comparisons, 4,794 unique observations remained with a total count of 186,984 observations. The mean number of observations for beta diversity comparison was 2,749±753.

Adequate sampling depth was assured by asymptotic approach of Chao1 rarefaction plot for all 3 timepoints (**Figure 4A**). Temporal effects were not sufficient to change phyla level abundance (**Figure 4B**), and did not cause Principal Component Analysis (PCOA) clustering (**Figure 4C**), suggesting that further subsampling is indicated to observe effects of medical school on the microbiome.

**Figure 4:**
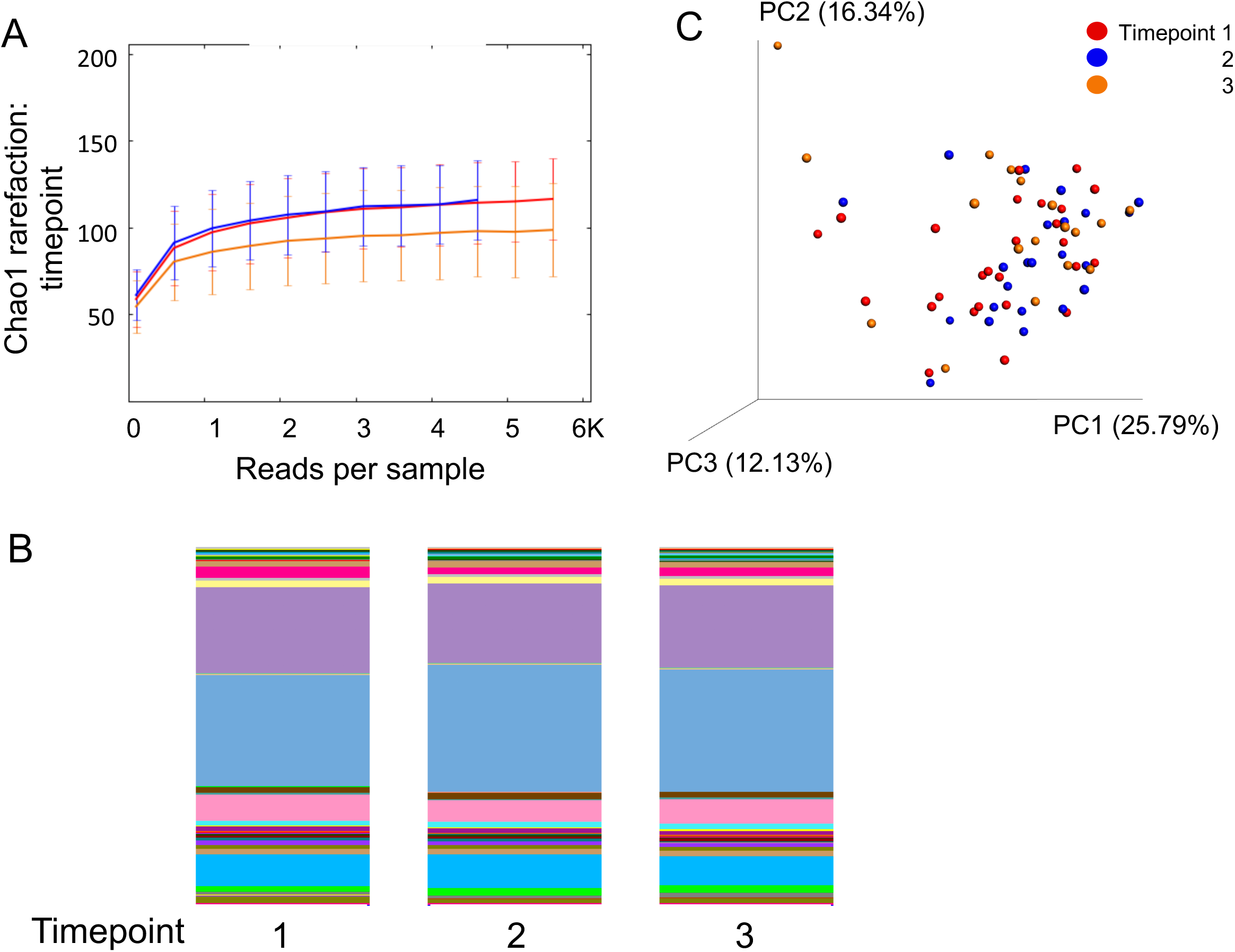
Temporal microbiome factors. Rarefaction of all timepoints shows that adequate sampling depth was reached at all timepoints (A). At the phyla level, no significant changes were observed across timepoints without subsampling the dataset (B). Principal Component Analysis (PCOA) found no groupings by timepoint (C).

### Phyla level relationship to stress resilience

Stress resilience subgrouping by identification of extreme resilience phenotype by RIS score stability showed more affects than temporal subsampling alone. As shown in **Figure 5**, at the OTU level, 6% of (294/4,795) observations were significantly different between the durable resilience and susceptible phenotypes before post-test adjustment by Kruskal-Wallis comparison. Of the 294 non-adjusted significantly different OTUs, 233 belonged to the Firmicute phyla, 36 from Bacteroidetes and 10 from Actinobacteria phyla. When these phyla level changes were observed between Firmicute and Bacteroidetes OTUs, phyla abundances were compared between groups. Of these observations, 2,102 OTUs were unique to the stress resilient phenotype on presence/absence comparison between groups. Despite the large proportion of microbiome differences between extreme resilience phenotypes, PCOA did not show a significant separation of groups.

**Figure 5:**
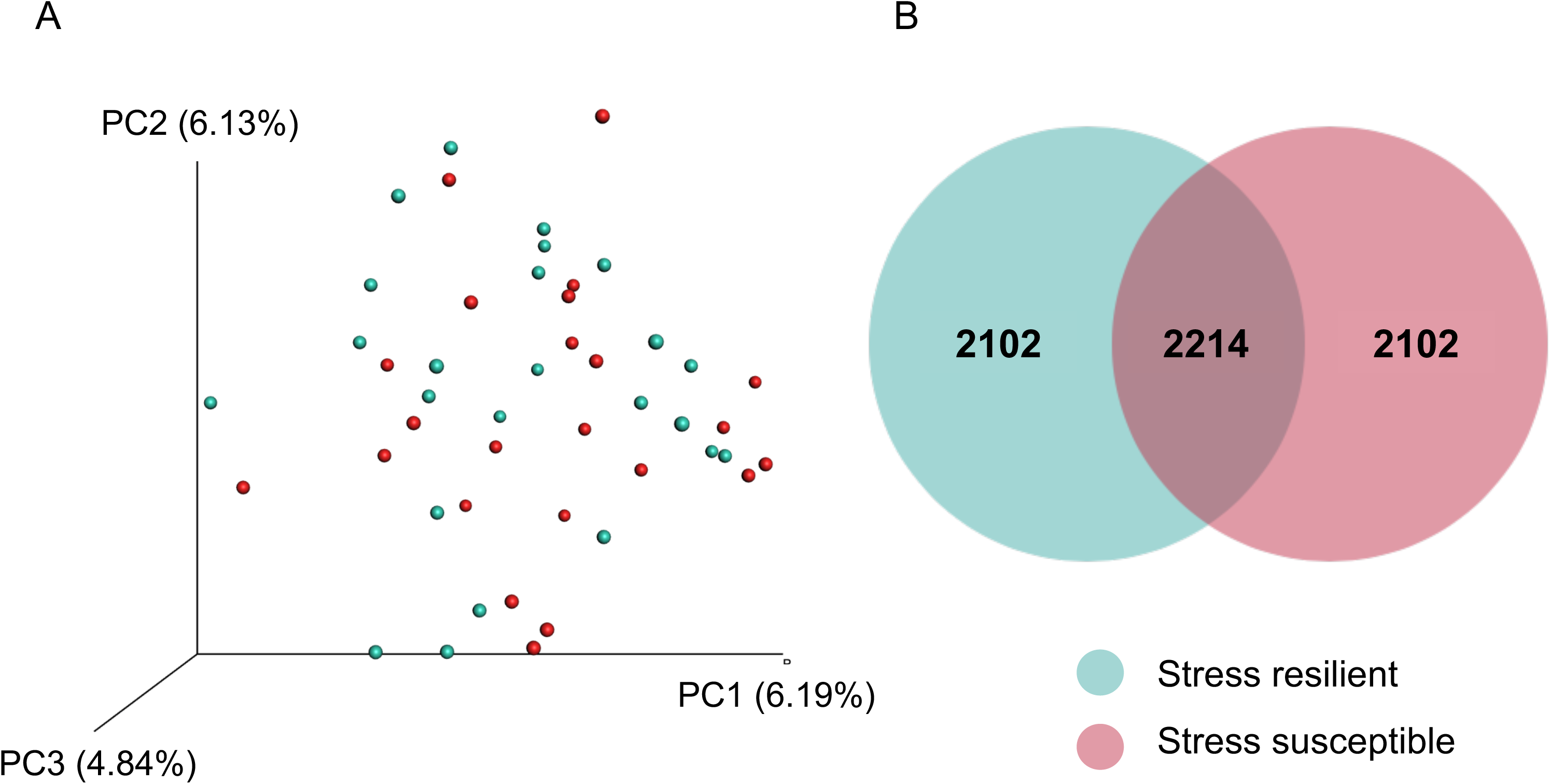
Evaluating microbiome changes during psychologic stress. **A)** Principal Component Analysis (PCOA) of microbiome beta diversity finds that the microbiome of durably resilient participants did not appear to be uniquely clustered. **B)** We found that 2,214 taxa were shared between groups, while 2,102 and 94 taxa were unique to the durable resilience and stress susceptible groups, respectively.

At the phyla level, no single phyla was significantly different between individuals with durable resilience. However, the Bacteroidetes:Firmicute ratio was found to have a significant negative correlation with RIS score (Pearson’s correlation, r=-0.24, p=0.05) (**Figure 6**).

**Figure 6:**
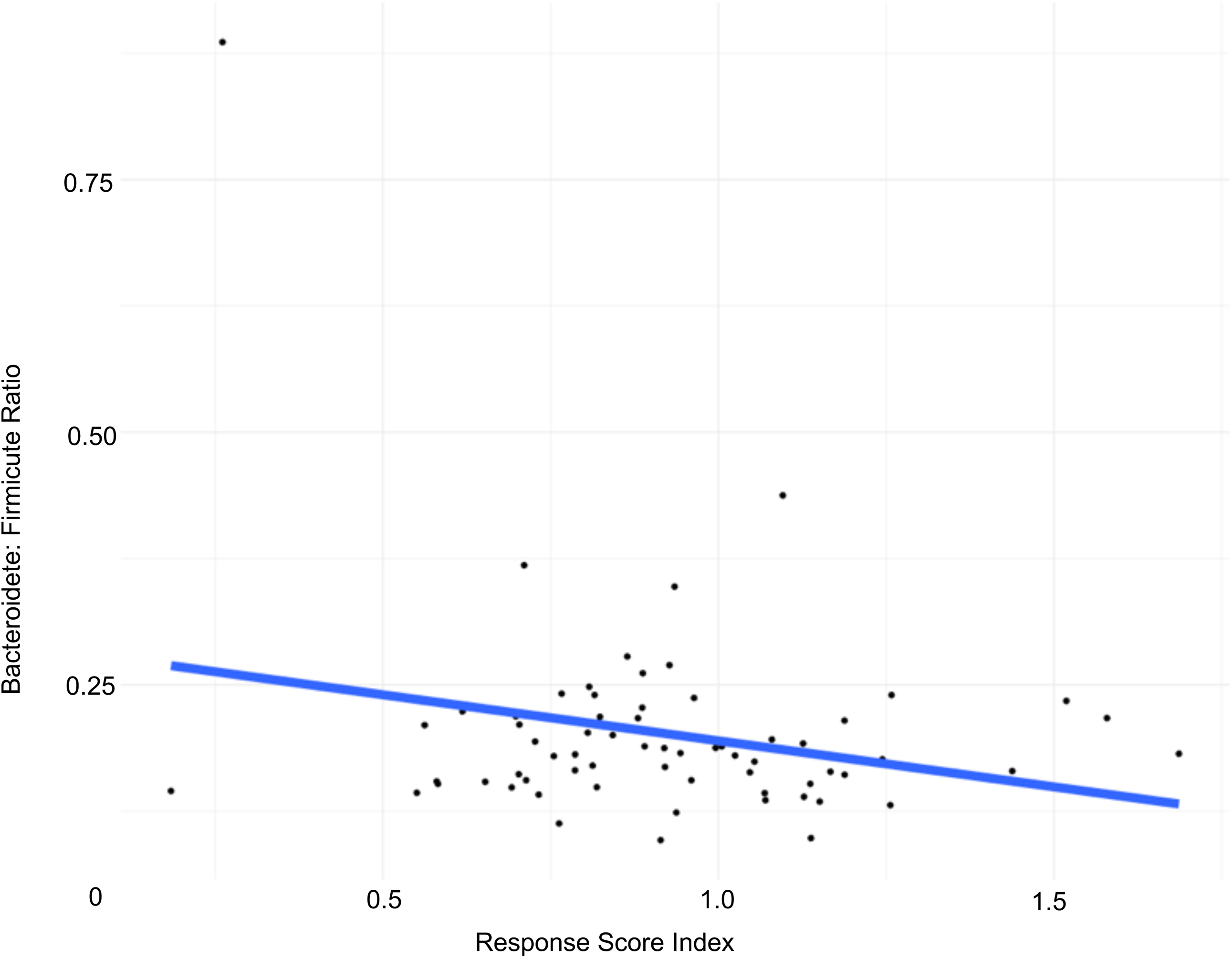
Bacteroidetes:Firmicute ratio. At the phyla level, the Bacteroidetes:Firmicute ratio displayed a significant negative correlation with RIS score (Pearson’s correlation, r=-0.24, p=0.05) in individuals with durable resilience.

### Assessment of temporal microbiome stability

Of 16 participants for whom we had 3 full timepoints of data available, temporal microbiome stability was found to be a feature of stress resilience. To assess temporal microbiome stability, a weighted Euclidean assembly of participants by microbiome relatedness was performed for participants with 3 timepoints of data (**Figure 7**). Participant microbiome stability was observed if all three timepoints of participant data clustered the participant’s microbiome samples into adjacent nodes. Microbiome instability was observed when participant microbiome samples did not cluster under adjacent nodes across all 3 timepoints. Temporal stability was observed in 13/16 participants. 8/8 participants displaying a durable stress resilience phenotype exhibited temporal stability while only 5/8, 63% of the stress susceptible phenotype showed temporal microbiome stability.

**Figure 7:**
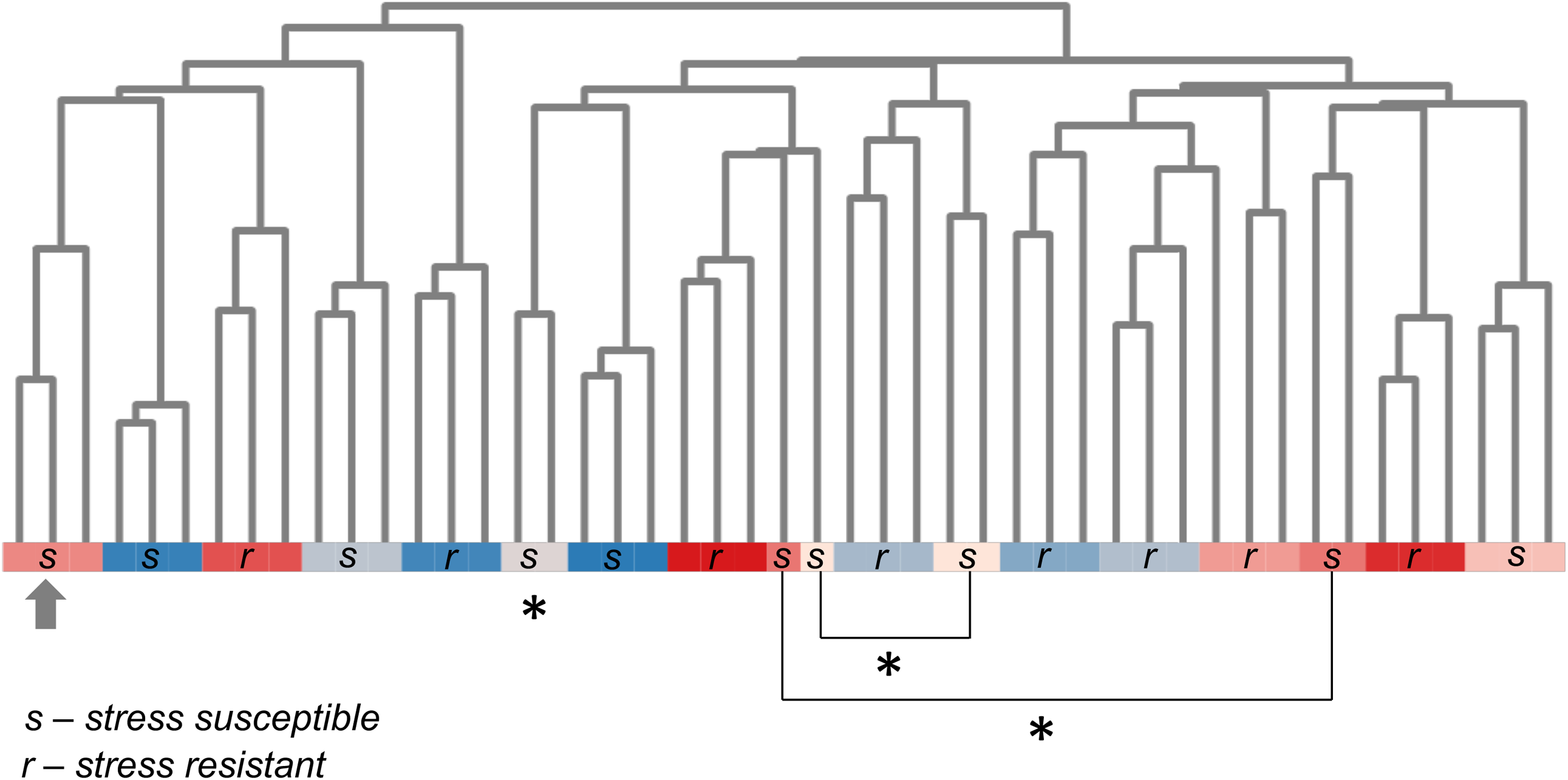
Temporal microbiome stability was observed in all participants with durable psychologic resilience. Weighted Euclidean assembly of 16 participants by microbiome relatedness was perfrmed for participants with 3 timepoints of data. The assembly was classified and colored by participant. Microbiome temporal stability was observed as when three timepoints of a participant cluster (arrow), and instability when three timepoints of a participant did not cluster adjacently (asterisks). 100% of durable resistance (r) exhibited sufficient temporal microbiome. Of the stress susceptible (s) group, 63% showed contiguous clustering.

## Discussion

### Microbiome stability and the regulation of stress perception

Prior work in the field of metagenomics has emphasized the role of microbiome stability on host physiology (Lozupone et al., 2012; Faith et al., 2013; Mehta et al., 2018; Tetel et al., 2018). What we found in this study was that phylogenetic assembly of longitudinal participant data by microbiome state may be an option to identify participants experiencing microbiome disruption events. There are two proposed mechanisms of how microbiome function may be a feature of stress resilience, in which the microbiome may act through bottom up or top down effects contributing to the observed stress phenotype (Foster et al., 2017; Tetel et al., 2018). In the top down hypothesis, the physiologic effects of stress changes the innervation of the gastrointestinal system, leading to a new physiological state and ultimately to novel niche selection within the gut lumen (Kelly et al., 2015). In this hypothesis, the microbiome plays a secondary role in which the now-altered transcriptome profile is a result of the host phenotype and further alters the perception of the stress phenotype. In the bottom up hypothesis, changes in gut lumen physiology alters gastrointestinal microbiome niche selection leading to a novel transcriptome which contributes to the host perception of the stress phenotype through production of neurotransmitters within the lumen (Rea et al., 2016; Foster et al., 2017). Because we were not able to quantify which change happened first, we are unable to conclude which hypothesis more accurately describes the nature of the role that the gut microbiome plays in human stress perception.

However, we did find support from our study that the microbiome does have a role in stress perception. We consider two possibilities that may be of interest to future work in this field. 1) Students experiencing increasing levels of stress throughout the school year had a greater magnitude of change in luminal physiology compared to their stress-resilient counterparts. Thus, there was greater top-down niche selection among stress susceptible students and that any microbiome changes are an incidental finding. An alternative theory is that 2) The magnitude of change to luminal physiology was constant between resilient and susceptible students when they were exposed to the same stressors, and that the differentiating feature was the ability of the participant microbiome to be change-resilient to novel niche selection. With microbiome disruption, as shown by phylogenetic dissimilarity, a feature of some stress-susceptible students we find support that further research into which mechanism is of greater importance in human resilience may be of benefit to students in stressful career paths.

### B:F ratio as a feature of stress perception

The Bacteriodetes:Firmicute ratio has been found as an important feature in body mass index, depression, ageing, and other physiological features (Ley et al., 2006; Mariat et al., 2009; Karl et al., 2018). What we found in this research was that the B:F ratio may also be a feature of stress resilience to chronic stressors. Owing to the complexity of the microbiome and the enormous number of features found within it, we were not expecting to find strong interactions between single components of the microbiome and host perception of stress. Rather we expected to find numerous moderate contributors of the microbiome to the perception of stress and found that one of these moderately important features was the Bacteroidete:Firmicute ratio. It will be interesting to observe whether this trend is represented by other studies using larger sample sizes.

### Medical students are an idealized model for stress resilience research

The study of medical student stress resilience is an area of opportunity for the future investigation of stress resilience among human subjects. In addition to the familiarity of IRBs to institutional research, medical student classes represent many other benefits which make them an ideal population for this area of study. First, they are an age matched cohort experiencing strong stressors simultaneously. This creates an opportunity to observe how many individuals of similar physical characteristics respond to consistent stress events. Among the benefit of temporally consistent stressors is the similar nature of the relocation event. Very few medical students within a class were not required to relocate for their education. This creates a ‘blank slate’ with a very similar starting point across a class, making an ideal opportunity for microbiome study of a population under stress as geographic effects on the microbiome are well known. By time-matching the relocation event and thus the common effect of new food sources, water and other environmental changes the microbiome effect of relocation becomes a common variable across the experimental population (Suzuki and Worobey, 2014). Future works may benefit from excluding students who did not relocate for medical school from their sample groups. This was not a concern for this study as Des Moines University draws students widely from the United States. Finally, it is well known that medical school is a powerful stressor which is taxing both mentally and physically to students making the study of intense chronic stress in a human population ethically permissible (Dyrbye et al., 2008; Hansell et al., 2019). A better understanding of the role of the gut microbiome in stress resilience in this particular population may raise the awareness of physician burnout and promote wellbeing of our future health care providers.

## Acknowledgements

We thank Zarin Rehman, Victoria Behrends, and Sarah Warywoda for their help with blood draws and sample organization. This work was supported by the Iowa Osteopathic Education and Research (IOER) Foundation.

## Author Contributions

L. Y. and A. S. designed experiments; M. R. R., J. J. and C. J. carried out the experiments and analysis; M. R. R. and L. Y. wrote the paper.

